# Structural and practical identifiability of contrast transport models for DCE-MRI

**DOI:** 10.1101/2023.12.19.572294

**Authors:** Martina Conte, Ryan T. Woodall, Margarita Gutova, Bihong T. Chen, Mark S. Shiroishi, Christine E. Brown, Jennifer M. Munson, Russell C. Rockne

## Abstract

Compartment models are widely used to quantify blood flow and transport in dynamic contrast-enhanced magnetic resonance imaging. These models analyze the time course of the contrast agent concentration, providing diagnostic and prognostic value for many biological systems. Thus, ensuring accuracy and repeatability of the model parameter estimation is a fundamental concern. In this work, we analyze the structural and practical identifiability of a class of nested compartment models pervasively used in analysis of MRI data. We combine artificial and real data to study the role of noise in model parameter estimation. We observe that although all the models are structurally identifiable, practical identifiability strongly depends on the data characteristics. We analyze the impact of increasing data noise on parameter identifiability and show how the latter can be recovered with increased data quality. To complete the analysis, we show that the results do not depend on specific tissue characteristics or the type of enhancement patterns of contrast agent signal.

## 1 Introduction

Contrast agent transport models have been developed and used for decades in the analysis of concentration time course data derived from dynamic contrast-enhanced magnetic resonance imaging (DCE-MRI), and are essential elements of many computational biology approaches [1, 2, 3, 4, 5, 6]. DCE-MRI transport models are linear ordinary differential equations with compartments describing the concentration of the contrast agent (CA) in tissue and vasculature. These models assume the compartments are well-mixed, i.e., contrast agent distributes evenly throughout the compartment instantaneously, such that contrast concentration is only a function of time, and not space, within an individual voxel. Several models have been formulated [7] based on different assumptions and simplifications, describing the blood–tissue exchanges of the administered contrast agent between compartments. The kinetic parameters derived from these models have been applied to a wide range of biological settings, including cancer, and have provided compelling diagnostic and prognostic value [8] as well as detecting early response to treatment [9].

Here we consider the structural and practical identifiability of four nested contrast transport compartment models frequently used to analyze DCE-MRI data. Identifiability is a fundamental property to create models able to capture the dynamics shown in the data with well-determined parameters [10, 11]. The issue of model identifiability revolves around the question of whether it is possible to use data to accurately and uniquely estimate parameters in the model. Generally, the larger the number of compartments in a model, the higher the accuracy, but at the cost of higher analysis complexity. Moreover, the more parameters that are included, higher is the probability of having a non-unique combination of model parameters which can accurately characterize the dynamics, underpinning the concept of structural identifiability. Therefore, there is a need to investigate how the kinetic parameters involved in the class of nested models used in DCE-MRI analysis can be determined, and what affects their identifiability, even in simple cases. Model identifiability becomes especially important for biological systems due to limited availability and quality of the data available [12, 13, 14]. Moreover, the amount and the quality of the data have a strong impact on the parameter identifiability of some parameters and, thus, on model outcomes and predictions.

Structural identifiability determines if the model structure is well-defined, however it is not sufficient to ensure the robustness of model outcomes when dealing with experimental biological data to inform the model. This is where the study of practical identifiability takes importance and a careful analysis of both types of identifiability becomes vital. In this work, we analyze the entire family of nested contrast transport models from both structural and practical viewpoints. To The best of our knowledge, few works in the literature have analyzed both structural and practical identifiability aspects of transport models used for DCE-MRI data, mainly focusing on the practical identifiability and only for a subset of the models [15, 16, 17, 18, 19].

This work is organized as follows. In Section 2, we describe the theoretical aspects of structural and practical identifiability, DCE-MRI data, and the class of considered models. Next, in Section 3, we provide structural and practical identifiability analysis of the most complex of the family of models, the Leaky Tofts–Kety model, applied to glioblastoma brain cancer data. We comment on the outcomes and implication in Section 4. Finally, in the Supplementary Section S we provide results concerning the nested family of sub-models as well as additional breast cancer data.

## 2 Materials and Methods

### 2.1 Theory of nested DCE-MRI transport models

Compartmental models of contrast transport for DCE-MRI are constructed assuming that the total tissue CA concentration *C*_*t*_(*t*) can be modeled as the sum of the CA concentration in the plasma space (PS) (*C*_*p*_(*t*)) and in the extracellular extravascular space (EES) (*C*_*e*_(*t*)), i.e.

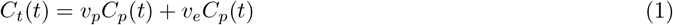

where *v*_*e*_ and *v*_*p*_ are the fractional EES and plasma volumes, respectively. The explicit description of the CA concentration in each of these compartments depends on the assumptions made to estimate both *C*_*e*_(t) and *C*_*p*_(t) and, thus, on the chosen model as illustrated in Figure 1A.

**Figure 1.**
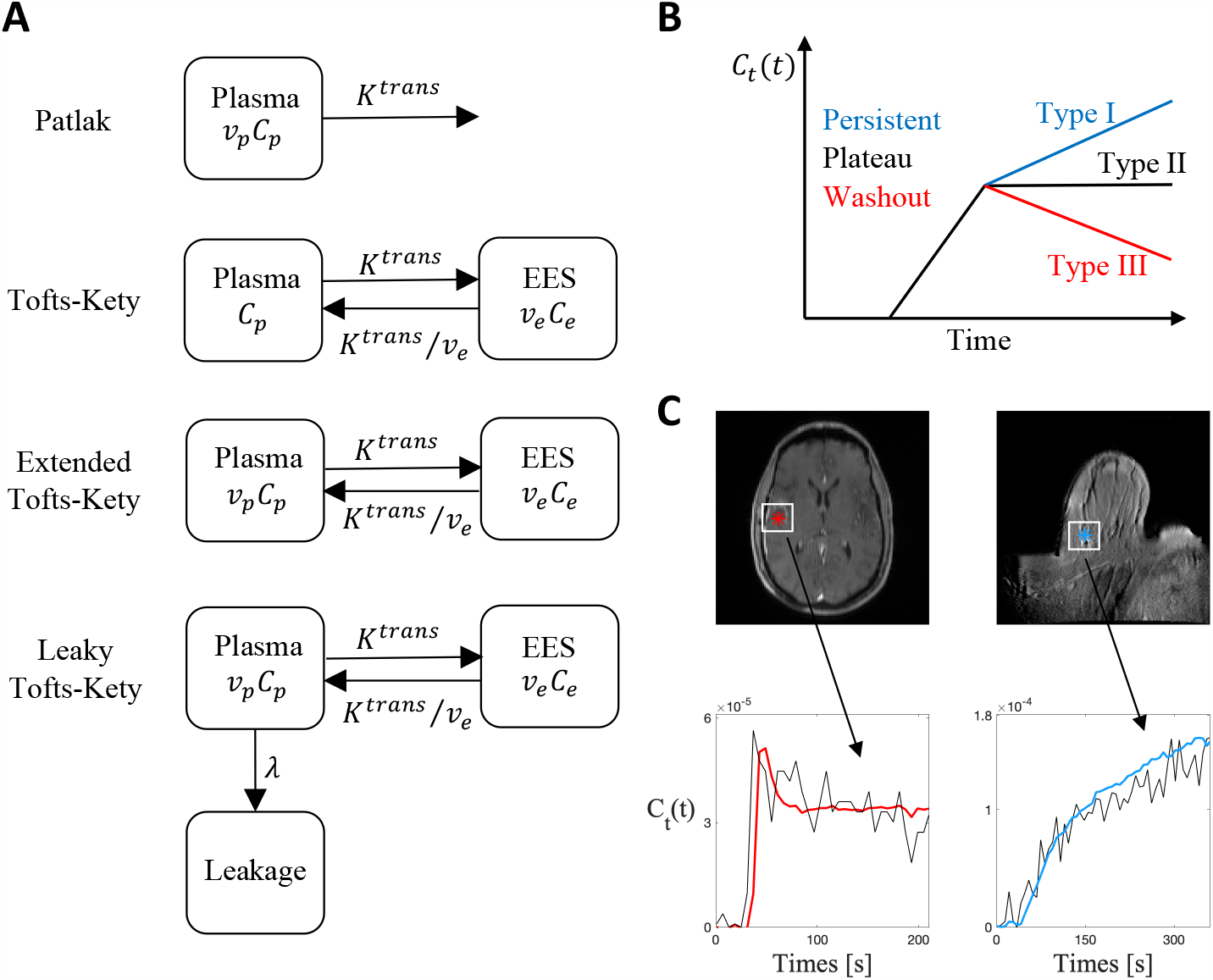
Models and data. **(A)** Schematic illustrations of the four nested transport models from simple (Patlak) to complex (Leaky Tofts–Kety): the contrast agent concentration *C*_*t*_(*t*) is evaluated using the functions *C*_*p*_(*t*), the CA concentration in the plasma compartment, which is assumed to be given by the arterial input function, and *C*_*e*_(*t*), for the CA concentration in the EES space. The rate of forward and backward volume transfer and the fractional EES and plasma volumes are the quantifies *K*^*trans*^, *v*_*e*_, *v*_*p*_, and λ. **(B)** Different enhancement patterns of CA signal: Type I is persistent, increasing contrast; Type II is a plateau following the initial peak; and Type III is washout with a decrease in contrast. **(C)** Examples of Type I (right plot) and Type III (left plot) enhancement curves obtained from breast and brain tumors, respectively.

The CA concentration for the plasma compartment, *C*_*p*_(t), is assumed to be not affected by local transport and it is given by the empirical vascular input function (VIF),

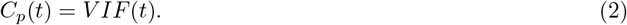

The dynamics of the VIF are characterized by a sharp uptake, followed by a peak value, and then a persistent, plateau, or washout dynamic (Figure 1B). Several techniques have been proposed to measure the VIF [20, 7]. Here we consider two different estimation methods. First, we consider a population-based analytical expression of the VIF, commonly referred to as the Parker VIF [21], and second, we estimate the VIF directly from the MRI signal [22]. The Parker VIF uses a mixture of 2 Gaussians plus an exponential modulated by a sigmoid function:

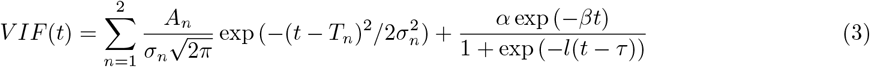

Here *A*_*n*_, *T*_*n*_, and *σ*_*n*_ are the scaling constants, centers, and widths of the n^*th*^ Gaussian, *α* and *β* are the amplitude and decay constant of the exponential, and l and *τ* are the width and center of the sigmoid, respectively. For the second method, VIF estimation is performed by drawing a region of interest (ROI) on a major vessel in the tissue and directly quantifying the contrast agent concentration, assuming the CA bolus arrives simultaneously to the entire tissue of interest. Population-based VIFs are widely used in DCE-MRI due to their simplicity and the fact that they do not require additional measurements, though they may ignore inter-subject variability. In contrast, the accuracy of individual-based VIFs derived from estimations of the CA concentration in large vessels depends on MRI characteristics, which may contribute to identifiability issues, as will become apparent through our analysis.

For the evolution of the CA concentration in the EES compartment, and, thus, to derive the expression for *C*_*t*_(*t*), we analyze a class of nested compartment models consisting of the Patlak model (PM), Tofts–Kety (TK) and its extended version (eTK), and the Leaky Tofts–Kety (LTK) model. The Patlak model, first introduced in [23], representing the simplest tissue uptake model, assumes that CA diffuses from the PS to the EES at rates governed by the forward transfer constant K^*trans*^, neglecting the reflux from the EES back into the plasma space due to low permeability and short measuring time. Thus, the concentration into the EES compartment varies according to

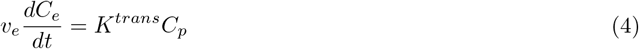

and, assuming that the initial concentration in the EES is zero (C_*e*_(0) = 0), from Eq. (1), the tissue CA concentration is given by

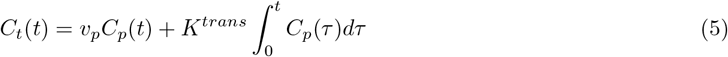

To overcome the low permeability hypothesis of the Patlak model the Tofts–Kety model was introduced in [24, 25]. The TK model accounts for bidirectional transfer of the contrast agent from the plasma to the EES compartment at rates governed by the forward transfer constant *K*^*trans*^ and the reverse constant *K*_*ep*_ = K^*trans*^/*v*_*e*_, respectively, however it ignores the intravascular (plasma) compartment contribution (i.e., *v*_*p*_ ≈0). The dynamics of the CA concentration in the EES compartment are described by

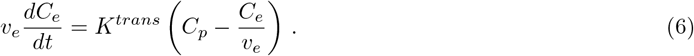

The transfer constant *K*^*trans*^ now has a different physiologic interpretation, depending on the balance between capillary permeability and blood flow: in high-permeability situations, *K*^*trans*^ is equal to the blood plasma flow per unit volume of tissue; in low permeability scenarios, *K*^*trans*^ is equal to the permeability surface area product between blood plasma and the EES, per unit volume of tissue [25]. Under the assumption that v_*p*_ ≈ 0 and setting the initial concentration in the EES to zero (*C*_*e*_(0) = 0), we can derive from Eq. (1) the expression of the tissue CA concentration as

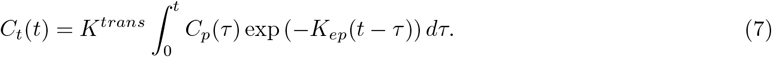

The hypothesis of tissue being weakly vascularized (*v*_*p*_≈ 0) may be invalid in some situations, especially in the case of highly vascular tumors. Thus, a modification of the TK model was proposed in [26]. Known as the extended Tofts–Kety model, it includes the vascular contribution *v*_*p*_*C*_*p*_(*t*) to the overall CA tissue concentration, maintaining the same dynamics shown in Eq. (6) for the EES concentration *C*_*e*_(*t*):

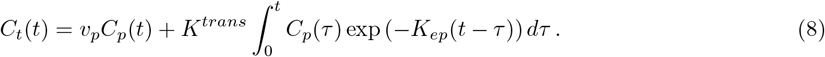

The TK and eTK models have experienced some problems to obtain reliable estimation for the involved parameters when the data depict a persistent uptake curve (Type I) [27]. The PM model, with its assumption on unidirectional flux from the plasma to the EES compartment, is able to overcome this issue, but assuming a negligible reflux from the EES back into the plasma space is not realistic, especially in the high permeability regime [28]. The Leaky Tofts–Kety (LTK) model was introduced to address this issue [27]. The LTK model adds an additional compartment, the leakage compartment, to the already described plasma and permeable compartments. The permeable space and the leakage space are considered subcompartments of the EES region. The former assumes a bidirectional exchange between plasma and EES, while the latter considers a unidirectional flow from which the contrast, at a concentration *C*_*L*_(*t*), does not flow back into the vasculature. Thus, for the LTK model, the overall CA tissue concentration is given by

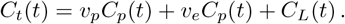

The contrast agent in the permeable compartment *C*_*e*_(t) evolves following Eq. (6), while the rate of contrast change in the leakage space can be read as

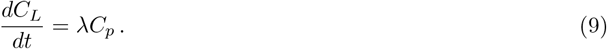

Here, λ is the volume transfer constant from the plasma compartment to the leakage compartment. Assuming that the initial concentration in the EES and leakage compartments are zero (*C*_*L*_(0) = *C*_*e*_(0) = 0), then the CA tissue evolution is given by

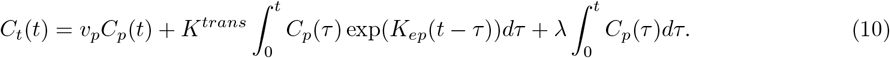

In summary, the main difference between these models are the directional flows for the CA exchange between the plasma space, the extravascular extracellular space, and eventually the leakage space. The Patlak model is a unidirectional model which assumes CA transfer from EES to PS is negligible due to low permeability and short measurement time [23]. The Tofts–Kety model assumes that the CA diffuses from, and returns to the PS, and the vascular (plasma) compartment contribution is negligible [24]. This assumption of tissue being weakly vascularized is overcome with the extended Tofts model, which includes the vascular contribution to the tissue concentration [26]. Finally, the Leaky Tofts–Kety model extends the eTK, adding an additional leakage compartment with unidirectional flow for disease applications such as gliomas, where slow accumulation of CA from neighbouring voxels is common [27]. In all models where the vascular component is considered, it is assumed that the concentration of the contrast agent within the vasculature is not affected by local transport, (i.e. is an empirical forcing function), and that the difference between the empirical vascular input function and the local tissue concentration is the primary driver of transport between tissue and vasculature.

### 2.2 Data

Two DCE-MRI scans were used in this study. A brain scan was obtained from a patient with pathology-confirmed diagnosis of glioblastoma who underwent MRI at City of Hope National Medical Center. The retrospective study was approved by the local institutional review board (IRB 15286) with a waiver of consent. The T1-weighted brain DCE-MRI scan was acquired as follows: TR=5.1ms, TE=2.1ms, variable flip angle, with field of view 240mm x 240mm, matrix size 128x128 and image size 256 x 256 with 12 slices with slice thickness 6mm, and a temporal resolution of 6.03 seconds with 32 dynamic phases and 3 baseline images prior to contrast administration. A breast DCE-MRI scan was obtained from the Quantitative Imaging Network (QIN) BREAST-02 study, UPN-01. The image acquisition details for the breast MRI can be found in the documentation for the BREAST-02 study [29, 30]. In both cases, a variable flip angle (VFA) scan was acquired for direct contrast agent quantification [31]. The VFA scan was used to calculate the baseline T1 relaxation, T_10_, of the tissue, which is used for calculating the local contrast agent concentration [32].

### 2.3 Identifiability analysis

Identifiability analysis is a fundamental mathematical tool used to assess a model’s capability to describe data with unique determined parameters. It focuses on whether it is possible to identify a unique vector of parameter values for a given model structure, or whether multiple parameter values will fit the data equally well. It is important to distinguish between *structural identifiability*, which concerns how the model structure itself affects the possible unique identification of the involve parameters, and *practical identifiability*, which is based on the analysis of the model’s ability to estimate parameters from the data with adequate precision.

Structural non-identifiability implies practical non-identifiability: in fact, if a model is not structurally identifiable, then the quality of the data collected does not matter; it will not be possible to uniquely estimate parameters in practice. Structural identifiability implies practical identifiability only when there is availability of an infinitely resolved data, with zero measurement noise. The converse, however, is not true; if the model structure theoretically allows parameters to be estimated, one still needs to have the appropriate data to achieve practically identifiable parameters.

#### 2.3.1 Theory for structural identifiability analysis

Consider a dynamical system

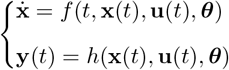

where **x**(*t*) ∈ ℝ^*n*^ represents the state variables, **y**(*t*) ∈ ℝ^*m*^ the measurable output (e.g., the data), h(·) the function that maps the states **x** to the observations **y, u**(*t*) ∈ ℝ^*r*^ the external input function (in our case the vascular input function), and ***θ*** ∈ ℝ^*q*^ the set of constant parameters. The dynamical system is structurally identifiable if each element *θ*_*i*_ of the vector ***θ*** is structurally identifiable. This means that each of these elements can be uniquely determined from a given input **u**(*t*) and a measurable output **y**(*t*), i.e.,

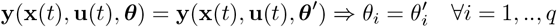

Alternatively, one element *θ*_*i*_ of the parameter vector ***θ*** is structurally non-identifiable if varying its value does not necessarily alter the model trajectory **y**, as these changes can be compensated for by varying other parameters. In particular, a model is defined to be structurally identifiable if all of its parameters are structurally identifiable [33]. A large variety of methods exist to assess structural identifiability of a system, from the Laplace transform [34], Taylor series expansion [35], seminumerical approaches [36, 37], differential algebra [38, 39, 40], and numerical algebraic geometry [41], among others (for reviews of some of these approaches, see Refs. [13, 42, 43]). Due to the simple structure of the nested models considered in this analysis described in Section 2.1, we apply a simple differential algebra approach. This approach is convenient for linear compartmental models because of the reduced number of state variables (i.e., *C*_*t*_(*t*)) and the direct relationship between state variables and observations (which actually coincides for this class of nested models). Further, this approach directly allows for testing identifiability, obtaining simple forms of possible identifiable combinations of the parameters, and identifying reparameterizations of the model when it results to be structurally non-identifiable.

#### 2.3.2 Theory of practical identifiability

As already briefly explained, a parameter that is structurally identifiable may still be practically non-identifiable. This issue arises frequently if the quantity and/or quality of the experimental data is insufficient. Assessing the practical identifiability of a model (or a parameter) is fundamental in obtaining reliable parameter estimations and, thus, for ensuring good model prediction capabilities. While the notion of practical identifiability is not uniquely defined in the literature, we consider a definition based on the concept of profile likelihoods and confidence intervals [44]. Using this framework, we evaluate the agreement between the experimental data and the observable predicted by our set of nested models choosing as Maximum Likelihood Estimator (MLE) the objective (or cost) function defined as:

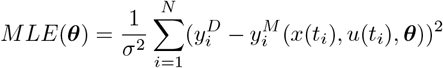

Here, *N* is the number of time points *t*_*i*_ available from the data, 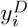 are the CA data values at time *t*_*i*_, 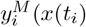, *u*(*t*_*i*_), ***θ***) is the CA value at time *t*_*i*_ predicted by the model with the estimation ***θ*** of the parameter vector, and *σ* is the standard deviation of the noise of the data. In particular, this standard deviation is estimated as the inverse of the Signal-To-Noise (SNR) ratio of the data[18]. The state variable *x*(*t*) represents the CA concentration *C*_*t*_(*t*) and *u*(*t*) is the vascular input function. The parameter vector 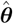 used to evaluate the practical identifiability of the model is given by

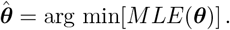

To determine the confidence interval of the estimated set of parameter 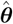, we consider the profile likelihood (PL). To build the profile likelihood plot, the idea is to choose one parameter *θ*_*i*_, vary its value around the maximum likelihood estimate 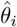, and re-optimize the remaining parameters. Thus, the profile likelihood is given by

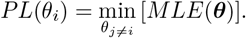

The profile likelihood method also estimates confidence intervals (CI) of estimated parameters defined as

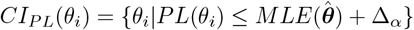

whereΔ_*α*_ represents the ff-th quantile of the chi squared (*χ*^2^) distribution with k = 1 degrees of freedom for a point-wise confidence interval [33, 45]. Thus, given the optimal value 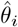 of one parameter, defining the confidence interval to a confidence level α implies that the true value 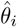 of the parameter is located within this interval with probability α. When this confidence interval is finite, than a parameter *θ*_*i*_ is practically identifiable. Otherwise, if the confidence region is infinitely extended, although the numerical algorithm finds a unique minimum 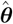, then the parameter is practically non-identifiable. Precisely, we analyze the profiles of the 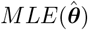 versus each parameter *θ*_*i*_.

A structurally non-identifiable parameter is characterized by a flat profile likelihood, whereas the profile likelihood of a practically non-identifiable parameter may have a minimum, but does not have a well-defined confidence level α for increasing and/or decreasing values of *θ*_*i*_. Instead, for an identifiable parameter the profile likelihood exceeds such threshold in both directions of *θ*_*i*_. Profile likelihood-based confidence intervals are often used in biological applications to ensure that there is sufficient data and data quality to assume the validity of the model [46, 47, 48, 11, 49]. With respect to other methods for determining confidence intervals, this particular method allows for asymmetric intervals that are invariant under re-parameterizations of the model.

In addition to the profile likelihood, we also analyze the compensating profiles of the model parameters. These are obtained by perturbing one parameter *θ*_*i*_ around its maximum likelihood estimate 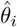 and plotting the variation of the re-optimized parameters versus the perturbed *θ*_*i*_ value. This analysis demonstrates how a non-identifiable parameter (whether structural or practical), may be compensated for by variations of other parameters. The small dimensionality of our systems allows us to use the described method for the analysis proposed in Section 3.2. We analyze the practical identifiability for the family of nested models described in Section 2.1 using a self-implemented code in Matlab based on the Particle Swarm algorithm for global optimization [50, 51].

## 3 Results

Here we analyze the results concerning structural and practical identifiability of the most complex of the nested models, the LTK model described in Section 2.1.

### 3.1 Structural identifiability of the LTK model

Following the formalism of the differential algebra approach described in Section 2.3.1, model (10) can be rewritten as

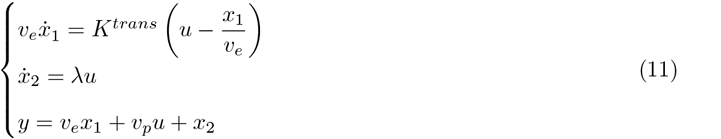

where *y* is the observable concentration of the contrast agent in the tissue *C*_*t*_(*t*), u(*t*) is the external input given by the concentration of contrast agent in the plasma compartment (i.e., *C*_*p*_(*t*) in (2)), *x*_1_ and *x*_2_ are the state variables for the CA concentration into the EES and leakage compartment, respectively. Differentiating the last equation in system (11) and plugging inside the first two equations of (11), we obtain the differential equation for *y*(*t*) of the form

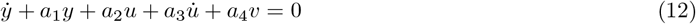

where *v* is defined as 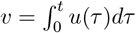 and the coefficients *a*_*i*_ are given by

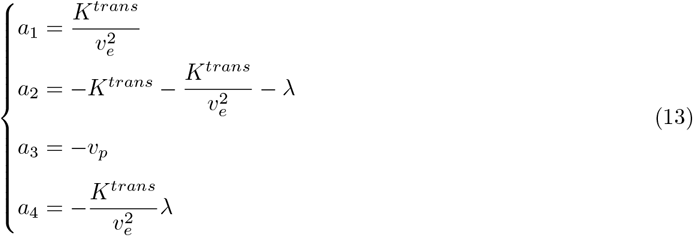

We observe that *a*_2_, *a*_3_, *a*_4_ < 0, while *a*_1_ > 0. Solving system (13) for the four parameters of the LTK model provides the following expressions:

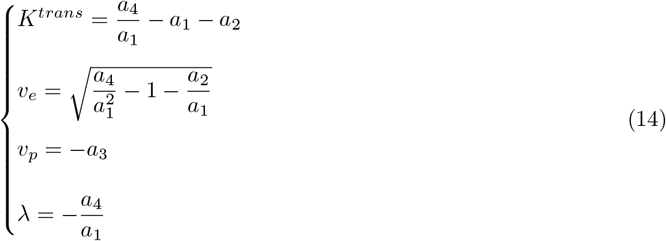

For *v*_*e*_ to be well-defined we need 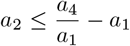, which always holds as the four parameters of the LTK model are positive. From this analysis we can conclude that LTK model is structurally identifiable. The structural identifiability of the nested PM, TK, and eTK models has also been analyzed and details are provided in the Supplementary Text S.1. Moreover, for the reader’s convenience, in the Supplementary Text S.2 we provide an example of a structurally non-identifiable model obtained by modifying the LTK model structure and changing the connections among the compartments.

### 3.2 Practical identifiability of the LTK model

Practical identifiability is performed using the method described in Section 2.3.2 and results are compared with the analytical results obtained in Section 3.1 regarding parameter structural identifiability. Precisely, we study the effect of varying the noise characteristics of the DCE-MRI data used for estimating the VIF and CA concentration on the accuracy of the identification of the model parameters. Each analysis is performed on the three characteristic enhancement patterns observed in CA evolution [7] (Figure 2). As shown in Figure 1B, CA time-enhancement curves are classified into three types: Type I - *persistent* curve — a progressive signal intensity increase (Figure 1C); Type II - *plateau* curve — is characterized by an initial peak followed by a relatively constant enhancement; Type III - *wash-out* curve — refers to a sharp uptake followed by an enhancement decrease over time (Figure 1C). We consider two different tissues to study the role of the tissue on model identifiability. In both tissues we replicate the analysis for the three different CA time-enhancement curves. We report results on the Glioblastoma multiforme (GBM) brain tumor data in the main text. Results for the breast cancer data are presented in the Supplementary Text S.3.1. For practical identifiability analysis we distinguish between the following three cases of study (referring the to the rows of Figure 2).

**Figure 2.**
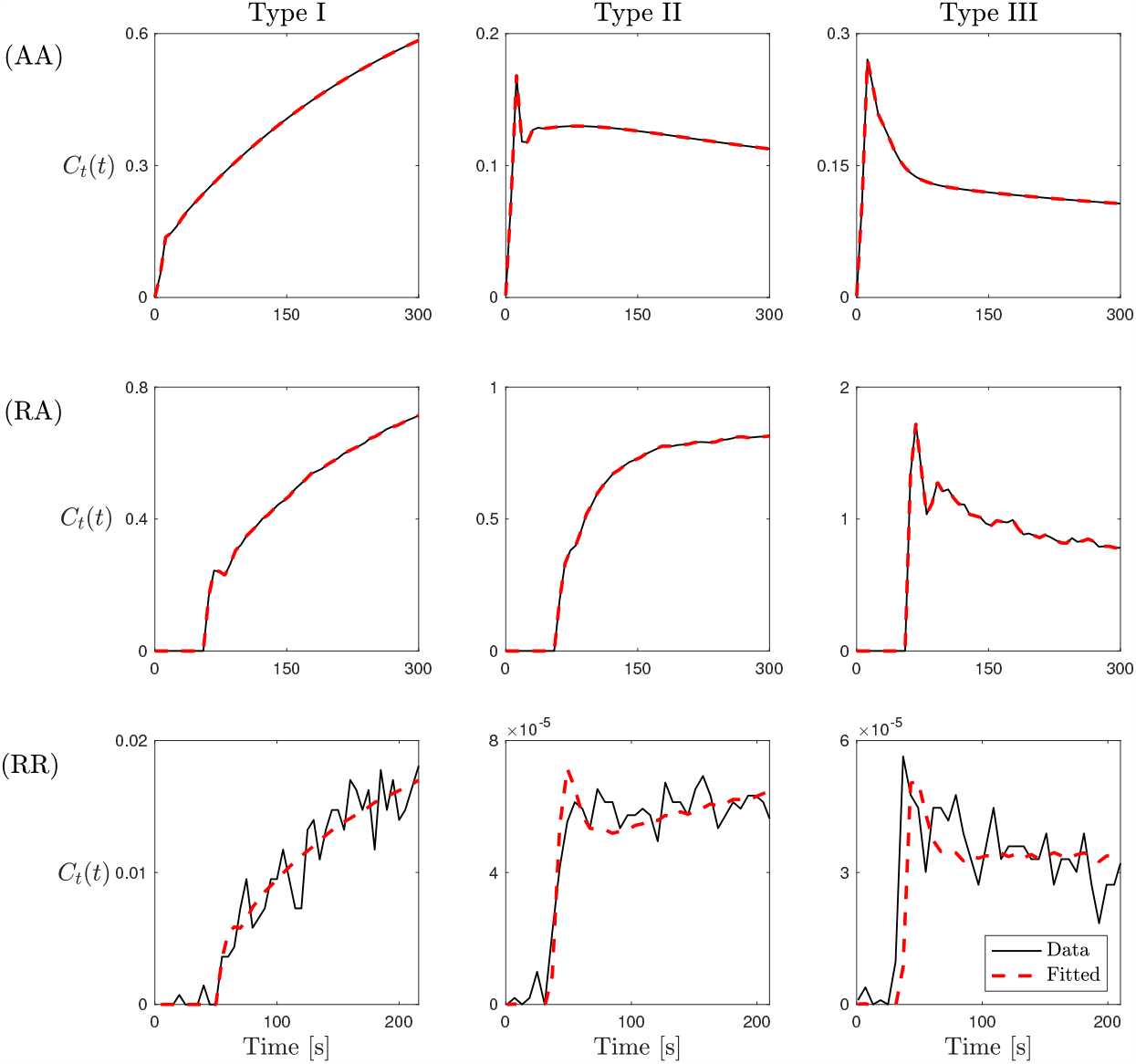
Best fitting of CA evolution signal obtained with the LTK model (10). The three types of CA time-enhancement curves (columns) for the three cases of study (AA), (RA), and (RR) (rows). The last row (RR) is VIF and CA time course data from the GBM DCE-MRI. Similar results for the breast cancer dataset are provided in Supplementary Figure S.6.

(AA) The first case is referred to as *Artificial-Artificial*. Here, we consider an *artificial* VIF given analytically by Eq. (3) and three *artificial* data sets computed using the LTK model (Eq. 9) to replicate the three CA time-enhancement curves. In this case, there is no noise in the VIF or in the data which could affect the parameter identifiability. For the VIF we set *A*_1_ = 48.54 mM × s, *A*_2_ = 18.64 mM × s, σ_1_ = 3.378s, σ_2_ = 7.92 s, *T*_1_ = 10.2276 s, *T*_2_ = 21.9 s, *α* = 1.05 mM, *β* = 0.0028 s^*-*1^, *l* = 0.6346 s^*-*1^, and τ = 28.98 s. The LTK model parameter values are summarized in Table 1.

**Table 1:** Parameter values used for the synthetic data set in the (AA) case.

| Type | $K^{trans}$ ($s^{-1}$) | $v_e$ | $v_p$ | $\lambda$ ($s^{-1}$) |
| --- | --- | --- | --- | --- |
| I | 0.0025 | 0.9 | 0.05 | 0.01 |
| II | 0.009 | 0.5 | 0.1 | 0.001 |
| III | 0.03 | 0.5 | 0.1 | 0.001 |

(RA) The second case is referred to as *Real-Artificial*. Here, we consider a *real* estimation of the VIF directly from DCE-MRI data by drawing a ROI around a major vessel, and three *artificial* data sets. The *real* VIF includes noise, as compared to the analytic expression for the *artificial* VIF as in the (AA) case. The *real* VIF and three *artificial* data sets are obtained with the parameter values summarized in Table 2.

**Table 2:** Parameter values used for the synthetic data set in the (RA) case of study.

| Type | $K^{trans}$ ($s^{-1}$) | $v_e$ | $v_p$ | $\lambda$ ($s^{-1}$) |
| --- | --- | --- | --- | --- |
| I | 0.0025 | 0.9 | 0.05 | 0.001 |
| II | 0.009 | 0.5 | 0.1 | 0.001 |
| III | 0.008 | 0.001 | 0.9 | 0 |

(RR) The last case is referred to as *Real-Real*. Here, we consider both a *real* estimation of the VIF, and a *real* evolution of the contrast agent in the tissue. We analyze data from a GBM brain tumor and from a breast cancer MRI dataset.

The three CA time-enhancement curves used in (AA), (RA), and (RR) case are shown in Figure 2 (continuous black lines) together with the corresponding model fits obtained with the LTK model (10) (dashed red lines). The particle swarm (PS) method is used to estimate parameter values for fitting the three curve type and in the three cases of study (AA), (RA), and (RR). Values of fitted parameters are collected in Supplementary Table S.1.

Recalling the methodology described in Section 2.3.2, we analyze profile likelihood and confidence levels for each of the parameters involved in the LTK model, in the three cases of study (AA), (RA), and (RR), and for the three types of CA time-enhancement curves. Here, we report the results for the Type I enhancement curve, as the LTK model was introduced with the aim of solving the issues of PM, TK, and eTK models concerning parameter estimation for persistent uptake CA profile. Results concerning Type II and Type III time-enhancement curves are collected in Supplementary Figures S.4 and S.5. Moreover, we show the results for *K*^*trans*^ and λ parameters. *K*^*trans*^ is the only parameter appearing in all nested models analyzed in this study, thereby allowing for a direct comparison of the results obtainable with the different models. λ is a novel parameter from the LTK model, thus it is important to analyze its identifiability.

Figure 3 shows the results concerning practical identifiability of the parameter *K*^*trans*^ for the LTK model and Type I time-enhancement curve.

**Figure 3.**
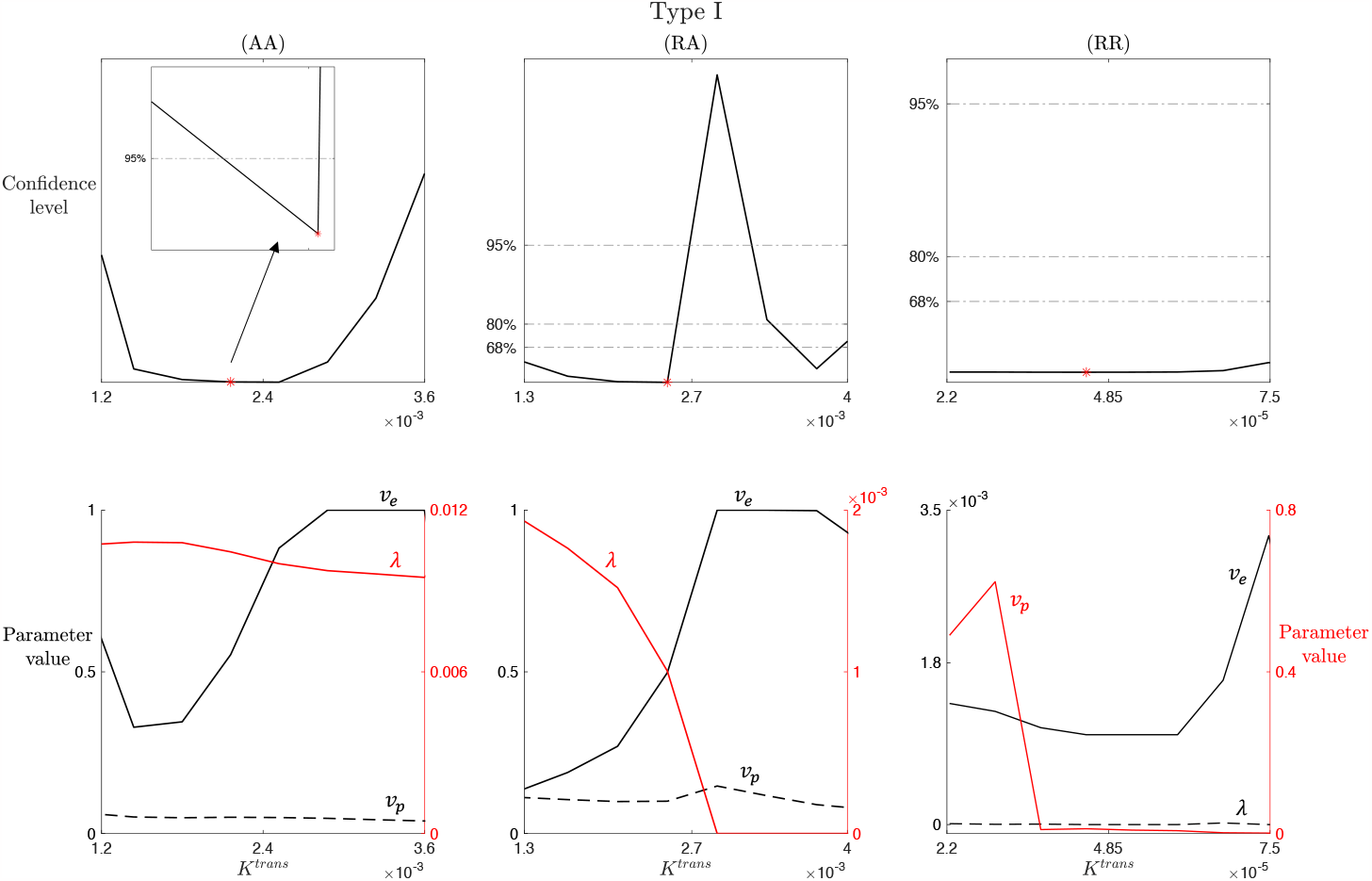
*K*^*trans*^ practical identifiability for LTK model and Type I enhancement curve. Top row: profile likelihood and confidence levels at 68%, 80%, and 95% for the parameter *K*^*trans*^ in the (AA), (RA), and (RR) case for the Type I enhancement curve. Inset in the first subplot shows a zoom of the region around the best fitted value 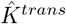 (red marker). Bottom row: compensating profiles of the parameters *v*_*e*_, *v*_*p*_ and *>* with respect to variation of *K*^*trans*^ around its best fitted value. Variation of *±*50% around the optimal value of *K*^*trans*^ are considered.

The columns of Figure 3 refer to the three cases of study (AA), (RA), and (RR). From Section 3.1, we know that *K*^*trans*^ is a structurally identifiable parameter, i.e., it is possible to uniquely determine its value from the given model structure. When we consider a completely artificial data set, obtained with the parameter values collected in Table 1 and the population-based analytical expression of the VIF given in (3), the complete absence of noise in the data and in the VIF allows us to accurately recover *K*^*trans*^. In the (AA) case, the profile likelihood shows a well-defined parabola, with a unique minimum in the optimal value 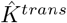. Zooming around this minimum, we observe a finite confidence region around 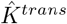 for the 95% confidence level. Analyzing the (RA) case (second column), we notice that 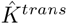 represents a local minimum for the profile likelihood and it is not possible to identify a lower bound for the confidence regions for both 95%, 80%, and 68%, i.e., it is not possible to define a finite confidence interval. Thus, *K*^*trans*^ is practically non-identifiable in this case. Moreover, the compensating profiles in the region where the profile likelihood is almost flat (left side with respect to 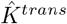, we notice *v*_*e*_ and λ compensating for varying *K*^*trans*^. In contrast, *v*_*p*_ is not significantly influenced by *K*^*trans*^ variation. In the (RR) case, *K*^*trans*^ is not practically identifiable, evident from the flat likelihood profile, which makes it not possible to identify a lower or upper bound of a confidence region with respect to the optimal value 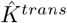.

Similar observations can be made from the results of Figure 4, where practical identifiability of the parameter λ for the LTK model and Type I time-enhancement curve are analyzed. As for Figure 3, columns of Figure 4 refers to the three cases (AA), (RA), and (RR). From the structural identifiability analysis of Section 3.1, we know that λ is a structurally identifiable parameter, thus, it is possible to uniquely identify its value from the model structure. For completely artificial data set used in (AA), we are able to accurately identify λ. The profile likelihood in the first subplot of Figure 4 shows a well-defined parabola with a unique minimum in the optimal value 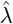, around which a finite confidence region for the 95% confidence level can be determined (as shown in the zoomed inset). Analyzing the RA and RR cases (second and third columns), we notice that the profile likelihood is characterized by flat regions for which it is not possible to identify a lower or upper bound for 95%, 80%, and 68% confidence levels. Thus, λis not practically identifiable in these more realistic cases. Looking at the corresponding compensating profiles, we observe that *v*_*p*_ estimation is not influenced by λ variation, while *v*_*e*_ and *K*^*trans*^ show variability in response to λ. This result is reasonable in relation to the results in (14): the expression for v_*p*_ is independent from the other parameters, whereas *K*^*trans*^, *λ*_*e*_, and λ expressions depends on common coefficients (i.e., a_1_, a_2_, and a_4_).

**Figure 4.**
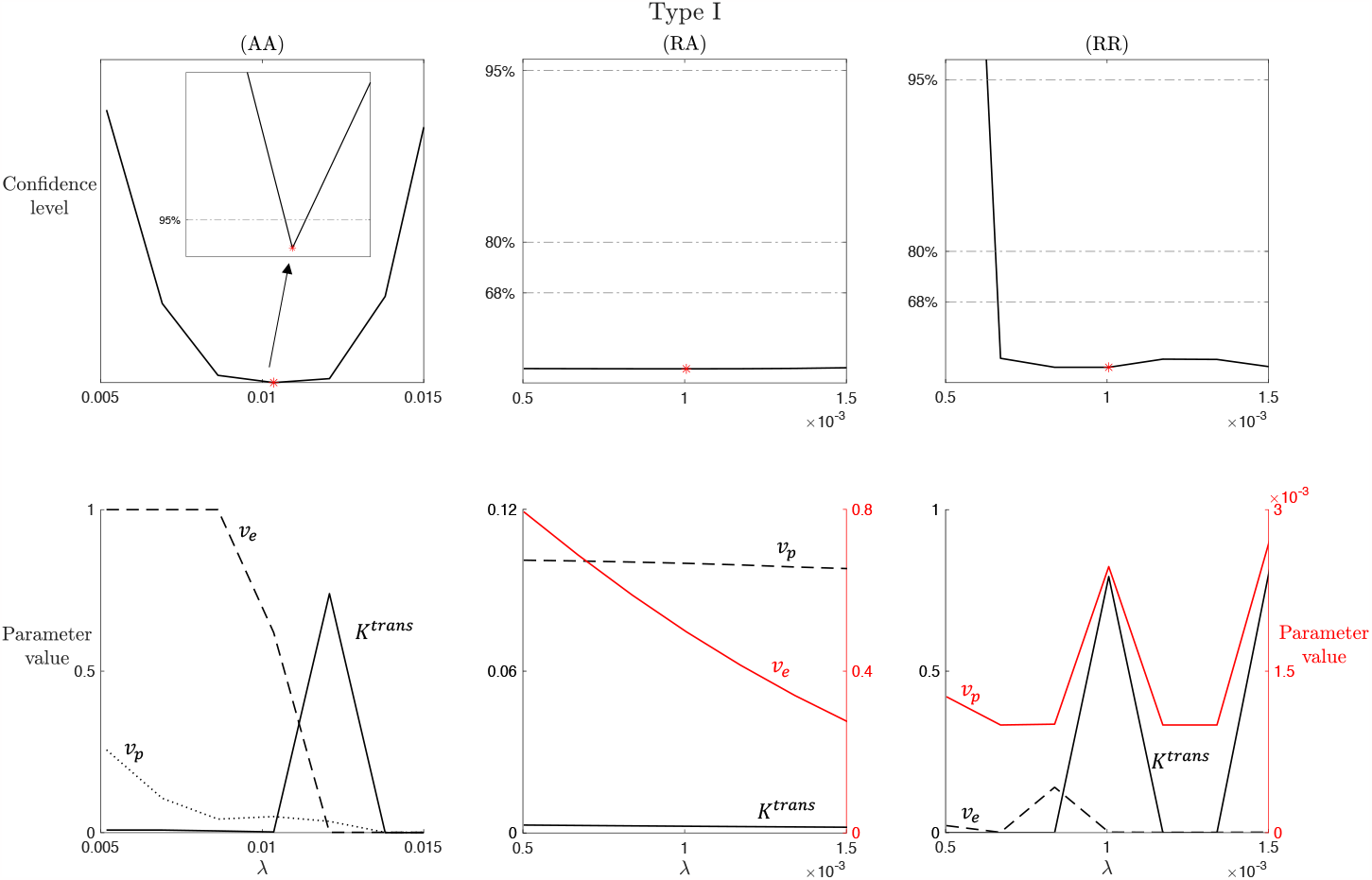
Leakage (*>*) practical identifiability for LTK model and Type I enhancement curve. Top row: profile likelihood and confidence levels at 68%, 80%, and 95% for the parameter *K>* in the (AA), (RA), and (RR) case for the Type I enhancement curve. Inset in the first subplot shows a zoom of the region around the best fitted value 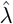 (red marker). Bottom row: compensating profiles of the parameters *K*^*trans*^, *v*_*e*_, and *v*_*p*_ with respect to variation of *>* around its best fitted value. Variation of *±*50% around the optimal value of *>* are considered.

To specifically analyze the influence of the noise in both the data and the individual-based estimation of the VIF function, derived directly from the DCE-MRI data, we perform two different tests, shown in Figures 5 and 6. We first analyze the impact of increasing the noise in the VIF function, mimicking the process that leads from the AA to the RA case. The individual-based estimations of the VIF are characterized by a predefined level of noise that is not present in the analytical VIF estimations. Figure 5 shows the results of this analysis for the parameter *K*^*trans*^ and Type I time-enhancement curve. Analogous results can be obtained with for the other parameters and enhancement profiles. The first column, the no noise case, replicates the same results shown in the first column of Figure 3, i.e., the practical identifiability of *K*^*trans*^ with a confidence level of 95%. Increasing the noise in the VIF, up to 5%, we observe that *K*^*trans*^ is still practically identifiable, but with a lower confidence level (80%). Considering higher levels of noise of the VIF (10% or 15%), the practical identifiability of the parameter is no longer guaranteed, as the confidence region bounds become not anymore finite for the considered confidence levels. This supports the conclusion that increasing noise in the VIF decreases the confidence level of parameter identifiability, justifying the differences in the results between (AA) and (RA) case in Figures 3 and 4.

**Figure 5.**
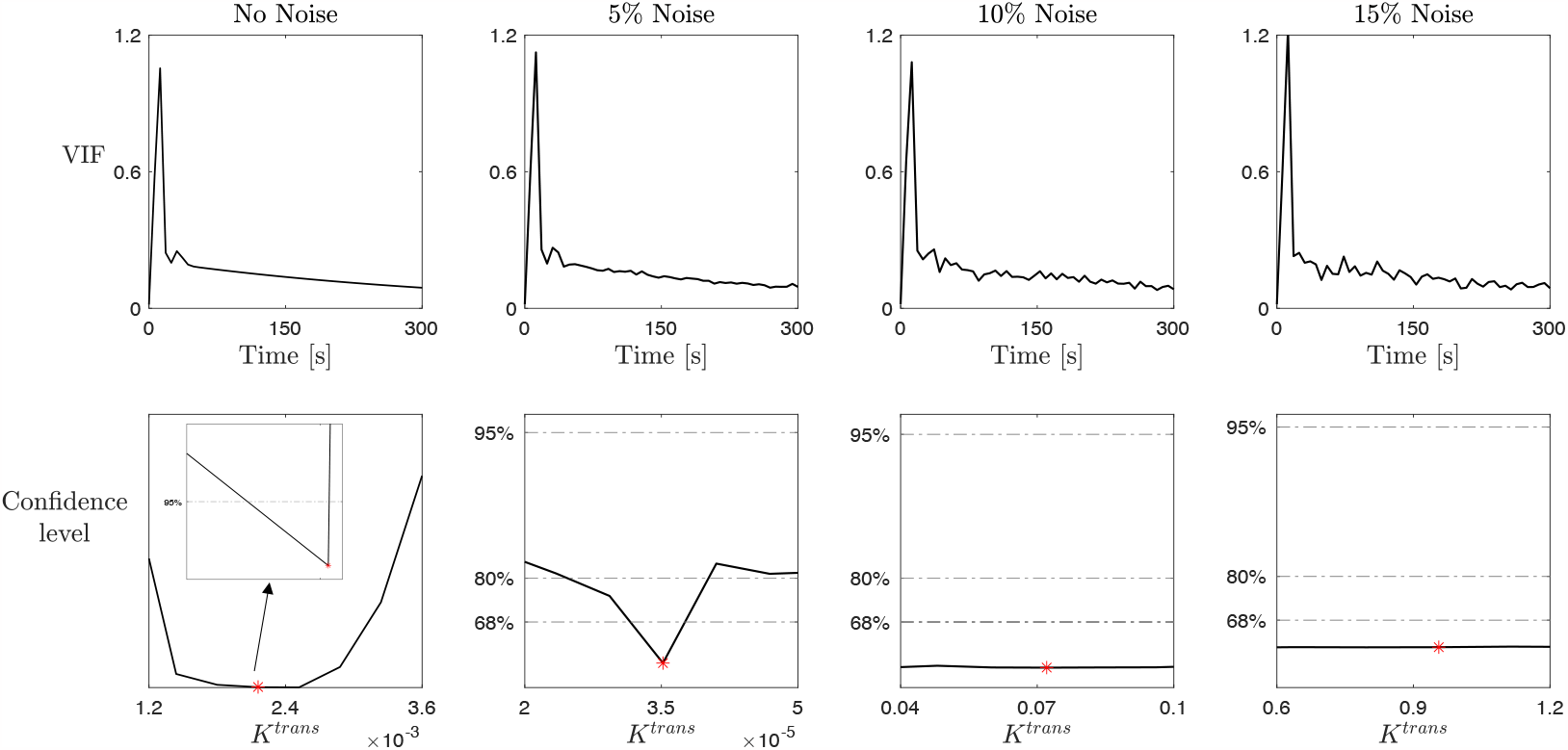
Study on the noise effect on parameter practical identifiability in the LTK model. Top row: Artificial VIF obtained from the analytical expression (Eq. (3)) by adding increasing amounts of noise. No noise VIF (first subplot) is the artificial VIF used in the (AA) case, while 5%, 10%, and 15% noise VIF (second to fourth subplots) are obtained by adding noise to the no noise case. Bottom row: profile likelihood and confidence levels at 68%, 80%, and 95% for the parameter *K*^*trans*^ for a Type I enhancement curve obtained by repeating the study on *K*^*trans*^ with the increasing noisy VIF illustrated in the top row. Red markers indicate the best value 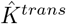.

**Figure 6.**
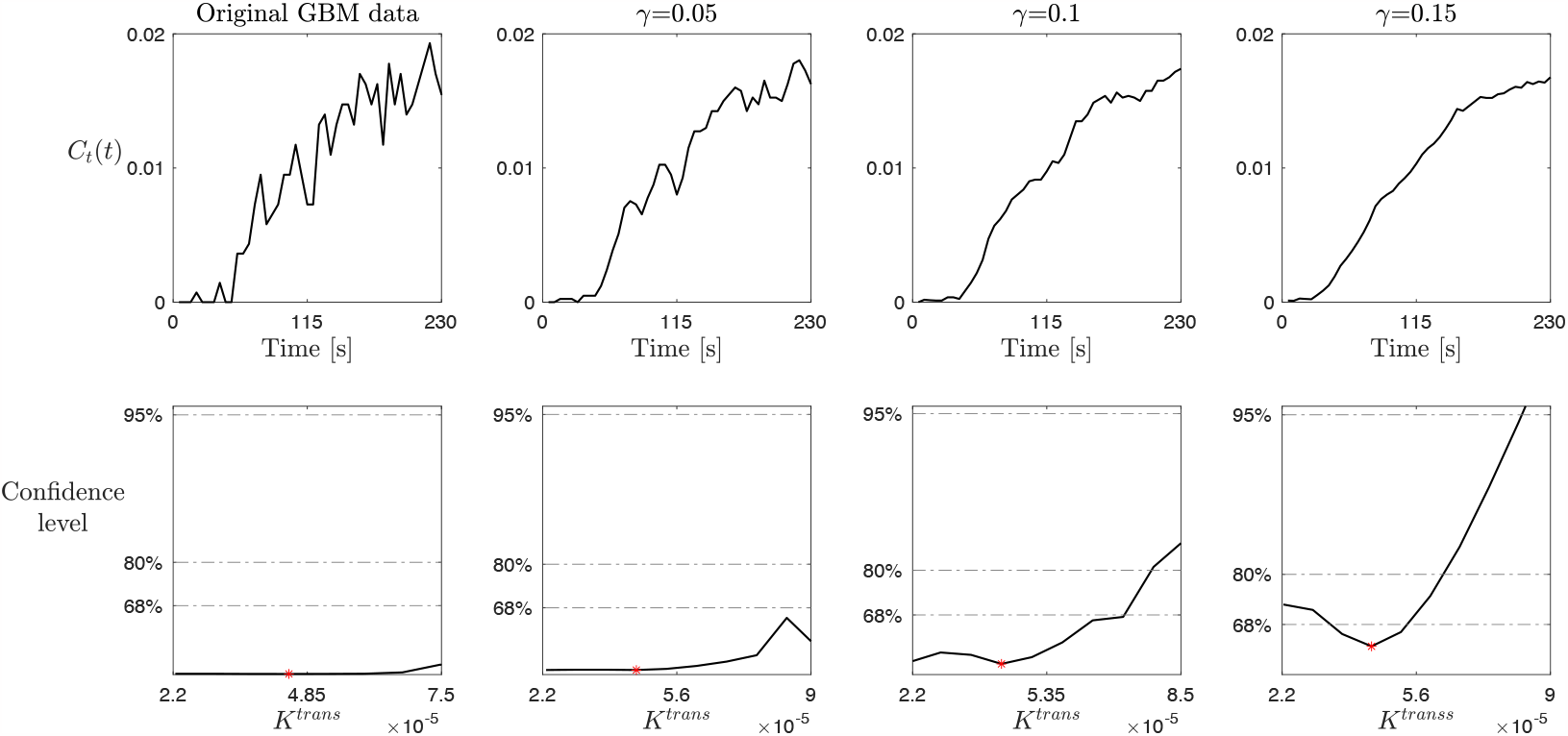
Study on the smoothing effect on parameter practical identifiability for the LTK model. Top row: CA profile obtain from the GBM data in a Type I enhancement curve by smoothing the data using a moving average method with different values of the smoothing factor γ*’*, i.e., γ*’* = 0.05 (second column), γ*’* = 0.1 (third column), and γ*’* = 0.15 (fourth column). The original GBM data (first column) are used in the (RR) case. Bottom row: profile likelihood and confidence levels at 68%, 80%, and 95% for the parameter *K*^*trans*^ obtained by repeating the study on *K*^*trans*^ with the smoothed data illustrated in the top row.

To analyze the influence of the noise in CA data on the parameter identifiability and, thus, mimicking the process that leads from the RR to the RA case, we perform a second test based on data smoothing. Starting from the CA time evolution data from the GBM dataset for a Type I time-enhancement curve, we use a moving average method with different value for the smoothing factor γ, which adjusts the level of smoothing by scaling the window size. Values of γ near 0 produce smaller moving window sizes, resulting in less smoothing, while values near 1 produce larger moving window sizes, resulting in more smoothing. Figure 6 summarizes the results of this analysis. The first column, referring to the original GBM data, replicate the same results shown in the last column of Figure 3, i.e., *K*^*trans*^ is not practically identifiable, with an almost flat profile likelihood. However, moving from left to right in Figure 6, the higher the value of the smoothing factor γ, i.e., the smoother the data, the less flat the corresponding profile likelihood. In particular, the last column of Figure 6, referring to γ = 0.15, shows the practical identifiability of *K*^*trans*^ can be recovered for the confidence level of 68%. We would not expect to obtain higher confidence level for *K*^*trans*^. These results are in line with the conclusion that high levels of noise in the data decrease the confidence level of parameter identifiability, in line with the results for RA and RR case in Figures 3 and 4.

## 4 Discussion

In this work, we have carried out a formal and in-depth study of the structural and practical identifiability of a well-known class of transport models widely used for the analysis of DCE-MRI data, with the aim of showing the impact of specific data characteristics on intrinsic features of these models. We focused on the LTK model, which accounts for the different aspects that are separately included in other members of the nested family of compartment models under investigation. However, analysis for the entire nested family has been also carried out and the corresponding results are shown in the Supplementary Information.

In Sections 3.1 and S.1.1, we analyzed structural model identifiability, showing how in all PM, TK, eTK, and LTK models it is possible to define a unique parametrization that identifies the parameters through the given inputs, i.e., CA concentration *C*_*t*_(*t*) and vascular input function VIF. As a matter of completeness, in Section S.2, we also described an additional transport model of the same class - namely the mLTK model - which is not structurally identifiable, but for which a structurally identifiable reparametrization can be defined. Then, in Section 3.2 and S.1.2, we used the method based on profile likelihood and confidence intervals to study practical identifiability of the nested family of models in three different scenarios, accounting for different levels of noise and construction of the data, and for different CA time-enhancement profiles. In fact, studying the practical identifiability of a model with respect to the data at hand is crucial for ensuring well-determined model predictions, as these are often used in systems biology research and quantitative image analysis.

The obtained results show how the entire class of transport models for DCE-MRI analysis from PM to LTK is structural identifiable, i.e., they have an intrinsic mathematical structure that allows for a unique identification of the parameters and, thus, reproducible and accurate outcomes. However, practically this would happen only in the ideal case of dealing with an large amount of data with zero noise. Our findings for both the LTK model (discussed in Section 3.2) and the nested PM, TK, and eTK models (discussed in Section S.1.2) reveal how practical parameter and, thus, model identifiability is affected by the quality of the data: the more noisy the data and/or the individual-based VIF function, the lower the confidence levels of identifiability of the different parameters. This is evident from the change in the shape of the profile likelihood transforming from a parabola-like shape in the (AA) case to the flat shape in the (RR) case, shown in Figures 3, 4, and S.1 for Type I contrast enhancement curve. The same change in the profile likelihood shapes are observed for Type II (Figure S.4) and Type III (Figure S.5) contrast enhancement curves. These results support the observation that model identifiability is attributed to a specific contrast agent dynamic, but rather on the characteristics of the data from which the enhancement curves are obtained.

The analysis proposed here is of significant importance considering the wide use of DCE-MRI data in research [52, 7] and, thus, the need for ensuring reliability and reproducibility of transport model results [53]. DCE-MRI has been shown to be associated with tumor angiogenesis and may be used to assess glioma grading [54, 55, 56, 57, 58, 59], predict genetic mutation status of brain tumors [60, 61, 62], distinguish pseudoprogression from true progression in glioblastomas [63, 64], and predict response to antiangiogenic treatment [65]. The parameter *K*^trans^, in fact, as a marker for tumor microvascular permeability from DCE analysis, could help to predict treatment response in glioblastoma [66]. Moreover, DCE-MRI parameters in a breast cancer study has been reported to be associated with microvessel density which is a marker for angiogenesis [67]. However, variability in DCE acquisition (e.g. in scan duration) and imaging analysis with various transport models may produce diverging results, which makes repeatability a challenging issue and hinders its wider adaptation for both clinical practice and research [53, 68]. Prospective longitudinal clinical trials, preferably in a multicenter setting with standardized imaging techniques and quantification of DCE parameters, are needed to validate the utility of transport parameters as imaging biomarkers for treatment response and survival prediction.

In this light, understanding model and data issues and limitations allows for a more conscious use of the results obtainable with them. Our results provide a mathematical explanation for the lack of repeatability of DCE-MRI quantification between clinical sites as has been previously reported [69]. As we have shown that practical identifiability improves with increasing SNR, we emphasize the need for image acquisition standards to increase the quality of imaging data and therefore reliability of parameter estimations such as the standards proposed by the Quantitative Imaging Biomarkers Alliance (QIBA) [70]. As DCE-MRI acquisition protocols and quantification methods continue to rapidly develop [71], it is important to ensure rigor in model fitting and analysis methods.

## Acknowledgement

Research reported in this publication was supported by the National Institutes of Health under award numbers P30CA033572, R01NS115971 (R.R., C.B., J.M.), R01CA254271 (C.B.) and the California Institute for Regenerative Medicine under award CLIN2-10248 (C.B.). The content is solely the responsibility of the authors and does not necessarily represent the official views of the National Institutes of Health or the California Institute of Regenerative Medicine. M.C. acknowledges also funding by the Ministry of Education, Universities and Research, through the MIUR grant Dipartimento di Eccellenza 2018-2022, project E11G18000350001, and the National Group of Mathematical Physics (GNFM-INdAM) through the INdAM–GNFM Project ”From kinetic to macroscopic models for tumor-immune system competition” (CUP E53C22001930001). The authors would like to thank the clinicians and researchers who contributed to the creation of the Quantitative Imaging Network Breast-02 and Treatment Response datasets, and the City of Hope neuro-oncology program. We especially thank all the patients who voluntarily participated in these studies, and their families, for their exemplary strength and generosity. We could not have performed this research without you.

## Author Contribution Statement

Conceptualization: M.C., R.W., R.R. Methodology: M.C., R.W., R.R. Formal analysis: M.C. Resources: R.R., C.B., J.M. Writing—original draft preparation: M.C. Writing—review and editing: M.C., R.W., M.G., C.B., J.M., B.C., M.S., R.R. Supervision: R.R. Project administration: R.R. Funding acquisition: C.B., J.M., R.R. All authors contributed to the article and approved the submitted version.

## List of Supplementary Information

**Supplementary Text S.1**. Identifiability of PM, TK and eTK models

**Supplementary Text S.2**. Modified Leaky Toft-Kety (mLTK): a structurally non-identifiable model

**Supplementary Text S.3**. Supplementary data for the LTK Model

**Supplementary Figure S.1**. *K*^*trans*^ **practical identifiability for PM (A), TK (B), and eTK (C) models and Type I enhancement curve**. Top row in **A, B**, and **C**: profile likelihood and confidence levels at 68%, 80%, and 95% for the parameter *K*^*trans*^ in the (AA), (RA), and (RR) case for the Type I enhancement curve. Inset in the first subplot shows a zoom of the region around the best fitted value of *K*^*trans*^ (red marker). Bottom row in **A, B**, and **C**: compensating profiles of the parameters *v*_*p*_ (in **A**), *v*_*e*_ (in **B**), and *v*_*e*_ and *v*_*p*_ (in **C**) with respect to variation of *K*^*trans*^ around its best fitted value. Variation of *±* 50% around the optimal value of *K*^*trans*^ are considered. The figure is embedded in the Supplementary Text S.1.

**Supplementary Figure S.2. Scheme for mLTK model** (S.4). Schematic illustration of the modified Leaky Tofts–Kety model: the contrast agent concentration *C*_*t*_(*t*) is evaluated using the functions *C*_*p*_(*t*), the CA concentration in the plasma compartment, which is assumed to be given by the arterial input function, and *C*_*e*_(*t*), for the CA concentration in the EES space. The rate of forward and backward volume transfer and the fractional EES and plasma volumes are the quantifies *K*^*trans*^, *v*_*e*_, *v*_*p*_, and λ. The figure is embedded in the Supplementary Text S.2.

**Supplementary Figure S.3**. *K*^*trans*^ **and leakage (**A**) practical identifiability for mLTK model and Type I enhancement curve**. Top row: profile likelihood and confidence levels at 68%, 80%, and 95% for the parameter *K*^*trans*^ (columns one to three) and A (columns four to six) in the (AA), (RA), and (RR) case for the Type I enhancement curve. Insets in the first and fourth subplot show a zoom of the region around the best fitted value 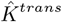 and 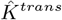 (red markers), respectively. Bottom row: compensating profiles of the parameters *v*_*e*_, *v*_*p*_, and λ with respect to variation of *K*^*trans*^ around its best fitted value (columns one to three) and of the parameters *K*^*trans*^, λ_*e*_, and *v*_*p*_ with respect to variation of λ around its best fitted value (columns four to six). Variation of *±*50% around the optimal values of *K*^*trans*^ and λ are considered.The figure is embedded in the Supplementary Text S.2.

**Supplementary Figure S.4**. *K*^*trans*^ **and leakage (**A**) practical identifiability for LTK model and Type II enhancement curve**. Top row: profile likelihood and confidence levels at 68%, 80%, and 95% for the parameter *K*^*trans*^ (columns one to three) and λ (columns four to six) in the (AA), (RA), and (RR) case for the Type II enhancement curve. Insets in the first and fourth subplot show a zoom of the region around the best fitted value 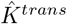 and 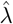 (red markers), respectively. Bottom row: compensating profiles of the parameters v_*e*_, v_*p*_, and λ with respect to variation of *K*^*trans*^ around its best fitted value (columns one to three) and of the parameters *K*^*trans*^, *v*_*e*_, and *v*_*p*_ with respect to variation of λ around its best fitted value (columns four to six). Variation of ±50% around the optimal value of *K*^*trans*^ and λ are considered. The figure is embedded in the Supplementary Text S.3.

**Supplementary Figure S.5**. *K*^*trans*^ **and leakage (**A**) practical identifiability for LTK model and Type III enhancement curve**. Top row: profile likelihood and confidence levels at 68%, 80%, and 95% for the parameter *K*^*trans*^ (columns one to three) and λ (columns four to six) in the (AA), (RA), and (RR) case for the Type III enhancement curve. Insets in the first and fourth subplot show a zoom of the region around the best fitted value 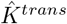 and 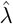 (red markers), respectively. Bottom row: compensating profiles of the parameters *v*_*e*_, *v*_*p*_, and λ with respect to variation of *K*^*trans*^ around its best fitted value (columns one to three) and of the parameter *K*^*trans*^, *v*_*e*_, and *v*_*p*_ with respect to variation of λ around its best fitted value (columns four to six). Variation of *±*50% around the optimal value of *K*^*trans*^ and λ are considered. The figure is embedded in the Supplementary Text S.3.

**Supplementary Figure S.6. Best fitting of the CA evolution with the LTK model** (9). The three types of CA time-enhancement curves (columns) are shown or the (RR) case of the breast cancer tissue. The figure is embedded in the Supplementary Text S.3.

**Supplementary Figure S.7**. *K*^*trans*^ **and leakage (**A**) practical identifiability for LTK model and Type I enhancement curve for breast cancer tissue**. Top row: profile likelihood and confidence levels at 68%, 80%, and 95% for the parameter *K*^*trans*^ (first column) and λ (second column) in the (RR) case for breast cancer tissue and for the Type I time-enhancement curve. Bottom row: compensating profiles of the parameters *v*_*e*_, *v*_*p*_, and λ with respect to variation of *K*^*trans*^ around its best fitted value (first column) and of the parameters *K*^*trans*^, *v*_*e*_, and *v*_*p*_ with respect to variation of λ around its best fitted value (second column). Red markers indicate the best fitted values 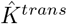 and 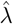. Variation of *±*50% around the optimal value of *K*^*trans*^ and λ are considered. The figure is embedded in the Supplementary Text S.3.

**Supplementary Table S.1**. The table is embedded in the Supplementary Text S.3

