## Supplementary Information for "Structural and practical identifiability of contrast transport models for DCE-MRI"

### S Supplementary Information

Here we present results which complete the analysis in the main text. In Section S.1 we report the results of structural (Section S.1.1) and practical (Section S.1.2) identifiability of Patlak, Tofts–Kety, and extended Tofts–Kety models. These belong to the same family of nested compartment models to which LTK belongs and their structure and derivation have been described in Section 2.1. In Section S.2 we illustrate an example of structurally non-identifiable model derived from a modification of the LTK model, showing the results about both structural and practical identifiability of its parameters. Finally, Section S.3 collects supplementary data that complete the analysis of the LTK model. In table S.1 we report the best parameter value obtained in the (RR) case together with Figures S.4 and S.5 for Type II and Type III time-enhancement curve for the parameters $K^{trans}$ and $\lambda$. Finally, results of the identifiability analysis of the LTK model in the case of breast cancer tissue are shown in Section S.3.1.

#### S.1 Identifiability of PM, TK and eTK models

In this Section, we analyze the structural and practical identifiability of the nested compartment models belonging to the same class of the LTK model and from which it has been derived. A complete description of their structure and related references have been provided in Section 2.1.

##### S.1.1 Structural identifiability for PM, TK, and eTK models

Assuming the leakage compartment and the reverse flux from extracellular space to the plasma space is negligible (i.e., $\lambda \approx 0$ and $C_e(t)$ evolution given by Eq. (4) in the manuscript), from the LTK model we reduce to the Patlak model. Using the formalism of the differential algebra approach introduced in Section 2.3, the PM can be written as

$$\begin{cases} v_e \dot{x} = K^{trans} u \\ \\ y = v_e x + v_p u \end{cases}$$

where $y$ is the observable concentration of the contrast agent in the tissue $C(t)$ and $u(t)$ is the external input given by the concentration of contrast agent in the plasma compartment (VIF). The state variable for the CA concentration into the EES compartment is given by $x$. Differentiating the second equation and combining it with the first, we obtain the differential equation for $y(t)$ in the form

$$\dot{y} + a_1 u + a_2 \dot{u} = 0$$

where

$$a_1 = -K^{trans} \quad \text{and} \quad a_2 = -v_p \,. \tag{S.1}$$

Thus, both parameters of the PM are structurally identifiable, allowing us to conclude that PM model is structurally identifiable.

The TK model can be obtained from the LTK by assuming that both leakage and intravascular compartment contributions are negligible (i.e., $v_p, \lambda \approx 0$). Using the same differential algebra approach, TK can be written as

$$\begin{cases} v_e \dot{x} = K^{trans} \left( u - \dfrac{x}{v_e} \right) \\ \\ y = v_e x \,. \end{cases}$$

This is analogous to the PM, as these two equations can be combined into the following equation for $y(t)$:

$$\dot{y} + a_1 y + a_2 u = 0$$

with

$$a_1 = \frac{K^{trans}}{v_e^2} \quad \text{and} \quad a_2 = -K^{trans} \,. \tag{S.2}$$

Thus, the TK model is structurally identifiable, as both $K^{trans}$ and $v_e$ are structurally identifiable.

Finally, the eTK model can be derived from LTK assuming that leakage compartment contribution is negligible (i.e., $\lambda \approx 0$). In the differential algebra formalism, eTK reads as

$$\begin{cases} v_e\dot{x} = K^{trans}\left(u - \dfrac{x}{v_e}\right) \\ y = v_e x + v_p u \end{cases}$$

whose equations can be recombined in the following expression for the evolution of the observable $y(t)$

$$\dot{y} + a_1 y + a_2 u + +a_3 \dot{u} = 0$$

with

$$\begin{cases} a_1 = K^{trans}/v_e^2 \\ a_2 = -K^{trans} \\ a_3 = -v_p\,. \end{cases} \tag{S.3}$$

Therefore eTK is also structurally identifiable, since all of its parameters are structurally identifiable.

##### S.1.2 Practical identifiability for PM, TK, and eTK models

Practical identifiability is performed using the method described in Section 2.3.2. As done for LTK model, each of these models has been analyzed in the three cases of study (AA), (RA), and (RR), described in 3.2 and for the three different time-enhancement profiles observed in CA evolution. For the purpose of this study and in analogy with LTK analysis, here we report the results only for the Type I time-enhancement curve and for the parameter $K^{trans}$. Figure S.1 collects the results concerning practical identifiability of this parameter for PM (S.1-**A**), TK (S.1-**B**), and eTK (S.1-**C**) models for Type I time-enhancement curve.

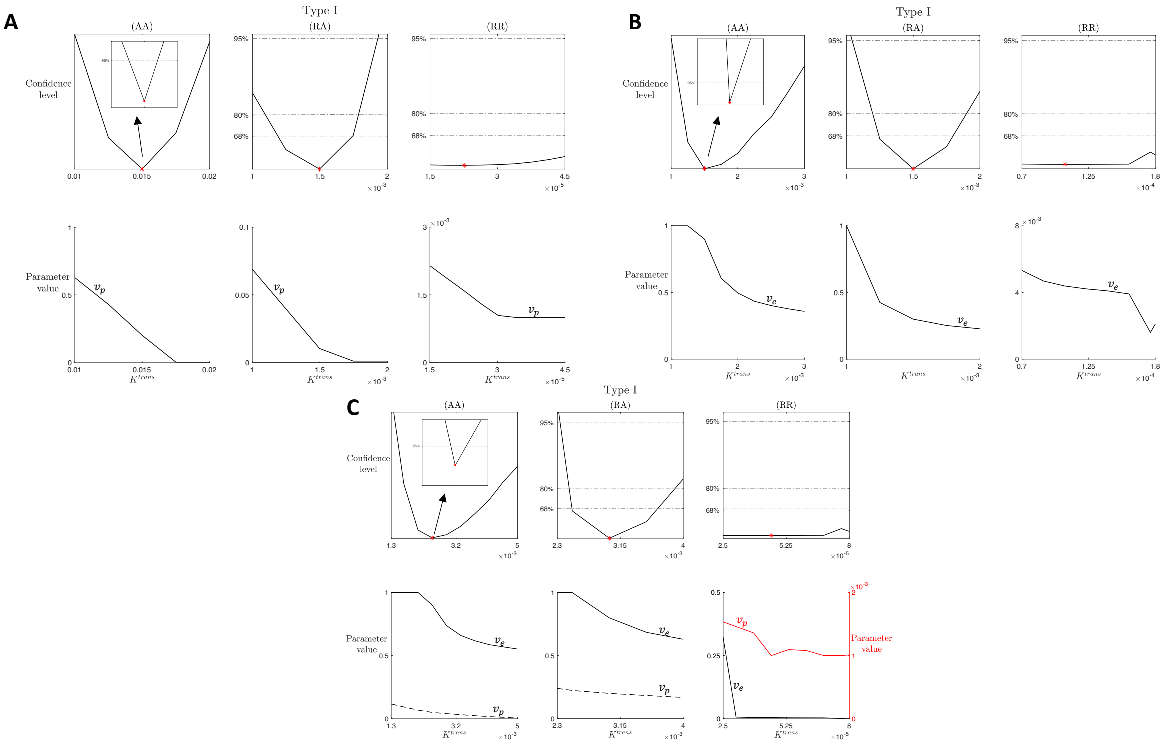


Figure S.1: $K^{trans}$ **practical identifiability for PM (A), TK (B), and eTK (C) models and Type I enhancement curve.** Top row in **A**, **B**, and **C**: profile likelihood and confidence levels at 68%, 80%, and 95% for the parameter $K^{trans}$ in the (AA), (RA), and (RR) case for the Type I enhancement curve. Inset in the first subplot shows a zoom of the region around the best fitted value of $K^{trans}$ (red marker). Bottom row in **A**, **B**, and **C**: compensating profiles of the parameters $v_p$ (in **A**), $v_e$ (in **B**), and $v_e$ and $v_p$ (in **C**) with respect to variation of $K^{trans}$ around its best fitted value. Variation of ±50% around the optimal value of $K^{trans}$ are considered.

The columns of each panel of Figure S.1 refers to the three cases of study (AA), (RA), and (RR), the top rows show the profile likelihood, and the bottom rows the compensating profile for the other parameters involved in the models. From the structural identifiability analysis of Section S.1.1, we know that $K^{trans}$ is structurally identifiable in PM, TK, and eTK, or equivalently, it is possible to uniquely obtain its expression from the given model structures. The columns referring to the (AA) case in **A**, **B**, and **C**, show that when an artificial data set is used, the practical identifiability of $K^{trans}$ is possible. In fact, the profile likelihoods show in all cases a well-defined parabola, with a unique minimum in the optimal value $\hat{K}^{trans}$ and the zoomed inset around this minimum shows a finite confidence region for the 95% confidence level. Analyzing the (RA) case (second columns in **A**, **B**, and **C**), instead, we notice that the confidence level for the identifiability of $K^{trans}$ in the three models decreases and we get a finite confidence region only for the 80% level, while it is not possible to define the lower/upper bound of the confidence region for 95% confidence level. This is in line with the results concerning the influence of VIF noise on parameter identifiability shown in Figure 5. Finally, looking at the (RR) results (third columns in **A**, **B**, and **C**), as for LTK model, we obtain flat evolution for the profile likelihood, meaning that the parameter $K^{trans}$ is practically non-identifiable. Compensating profiles (in the bottom rows of **A**, **B**, and **C**) show the response of the other model parameters to $K^{trans}$ variability around its best value. In particular, looking at eTK model (in **C**) we observe that $v_p$ has a rather small response to $K^{trans}$ variation, while $v_e$ seems to show more evident changes in response to $K^{trans}$. This result is reasonable in relation to what is shown in results in Eq. (S.3): in fact, the expression for $v_p$ is independent from the others.

#### S.2 Modified Leaky Toft-Kety (mLTK): a structurally non-identifiable model

In this Section, we illustrate an example of a structurally non-identifiable model for the analysis of DCE-MRI data. It belongs to the family of nested compartment models from which the proposed analysis is focused and it represents a modified version of the LTK model described in Section 2.1. An illustration of this mLTK model is shown in Figure S.2.

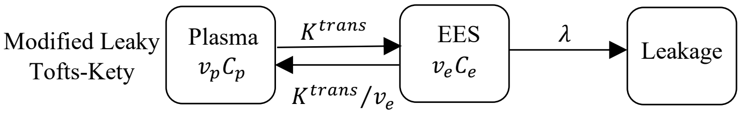


Figure S.2: **Scheme for mLTK model** (S.4)**.** Schematic illustration of the modified Leaky Tofts–Kety model: the contrast agent concentration $C_t(t)$ is evaluated using the functions $C_p(t)$, the CA concentration in the plasma compartment, which is assumed to be given by the arterial input function, and $C_e(t)$, for the CA concentration in the EES space. The rate of forward and backward volume transfer and the fractional EES and plasma volumes are the quantifies $K^{trans}$, $v_e$, $v_p$, and $\lambda$.

As with the LTK model, the mLTK includes a further compartment, the leakage compartment, which accounts for the loss of CA concentration that, coming out from the EES compartment does not flow back into the plasma compartment. Separately from the LTK, we do not consider a direct flow between PS and leakage compartment, but only a unidirectional flow from EES to the leakage compartment. Thus, the concentration of CA into the EES compartment varies according to

$$v_e \frac{dC_e}{dt} = K^{trans}\left(C_p - \frac{C_e}{v_e}\right) - \lambda C_e \tag{S.4}$$

and, assuming that the initial concentration of contrast agent in the EES is zero ($C_e(0) = 0$), from Eq. (1), the tissue CA concentration results in

$$C_t(t) = v_p C_p(t) + K^{trans} \int_0^t C_p(\tau) \exp\left(-\left(\lambda + K_{ep}\right)(t-\tau)\right) d\tau\,. \tag{S.5}$$

Using the formalism of the differential algebra approach introduced in Section 2.3, we can analyze the structural identifiability of the mLTK model. We rewrite Eq. (S.4) in the following form:

$$\begin{cases} v_e \dot{x} = K^{trans}\left(u - \dfrac{x}{v_e}\right) - \lambda x \\ y = v_e x + v_p u \end{cases} \tag{S.6}$$

where $y(t)$ is the observable concentration of the contrast agent in the tissue $C(t)$ and $u(t)$ is the external input given by the concentration of contrast agent in the plasma compartment (VIF). Differentiating the second equation and combining it with the first one, we can write the differential equation for $y(t)$ in the form

$$\dot{y} + a_1 y + a_2 u + a_3 \dot{u} = 0$$

where the coefficients $a_i$ are given by

$$\begin{cases} a_1 = \dfrac{1}{v_e}\left(\dfrac{K^{trans}}{v_e} + \lambda\right) \\ a_2 = -\left(K^{trans} + \dfrac{v_p}{v_e}\left(\dfrac{K^{trans}}{v_e} + \lambda\right)\right) \\ a_3 = -v_p \end{cases} \tag{S.7}$$

We observe that $a_2, a_3 < 0$, while $a_1 > 0$. Rearranging the terms in Eq. (S.7), we get

$$\begin{cases} K^{trans} = a_3 a_1 - a_2 \\ v_p = -a_3 \\ \lambda = a_2 - a_3 a_1 v_e + a_1 v_e^2 \end{cases} \tag{S.8}$$

From system (S.8) we notice that the parameters $K^{trans}$ and $v_p$ are structurally identifiable, while $\lambda$ and $v_e$ are structurally non-identifiable. The parameters $\lambda$ and $v_e$ are connected with the relationship

$$a_2 = \lambda + a_3 a_1 v_e - a_1 v_e^2$$

which has an infinite number or solution for $(v_e,\lambda)$. This suggests a structurally identifiable reparameterization of mLTK model. Defining a new parameter $\bar{K}^{trans} = \dfrac{K^{trans}}{\lambda}$ and a new time scale $\tau = t\lambda$, system (S.6) reads

$$\begin{cases} v_e \dot{x} = \bar{K}^{trans}\left(u - \dfrac{x}{v_e}\right) - x \\ y = v_e x + v_p u \end{cases} \tag{S.9}$$

where $\dot{x} := \dfrac{dx}{d\tau}$. Repeating the same differential algebra approached introduced above, from system (S.9) we obtain

$$\begin{cases} K^{trans} = a_1 a_3 - a_2 \\ v_e = \dfrac{1 + \sqrt{1 + 4a_1(a_1 a_3 - a_2)}}{2a_1} \\ v_p = -a_3\,, \end{cases}$$

i.e., the reparametrized mLTK model is structurally identifiable. Here, for $v_e$ to be well-defined we need $(a_1 a_3 - a_2) \geq 0$, which always holds as $(a_1 a_3 - a_2) = K^{trans}$ and the four parameters of the mLTK model have to be positive.

Concerning the practical identifiability of the mLTK model, we use the method described in Section 2.3.2, considering the three cases of study (AA), (RA), and (RR), described in 3.2, and the three different time-enhancement profiles observed in CA evolution. For the purpose of this study and in analogy with LTK analysis, we report the results for the Type I time-enhancement curve and for the parameters $K^{trans}$ and $\lambda$. Figure S.3 collects the results concerning practical identifiability of these parameters. Starting from the first three columns, referring to the parameter $K^{trans}$ in the (AA), (RA), and (RR) cases, we notice that the results are analogous to the ones shown in Figure 3. In fact, if the artificial data set used in (AA) allows to recover the practical identifiability of $K^{trans}$ (identified by the parabola-like profile with a unique minimum in the optimal value $\hat{K}^{trans}$ and a finite confidence region for the 95% confidence level), (RA) case shows a finite confidence region only for the 68% confidence level, while for (RR) case it is not possible to define the lower and upper bounds of the confidence region. Thus, the parameter $K^{trans}$ is practically non-identifiable, especially when real GBM data and individual-based estimation of the VIF are considered. Looking instead at columns four to six, we notice that none of the cases demonstrate identifiability for $\lambda$. In all cases, we observe that it is not possible to define the lower and upper bounds of the confidence region for all the considered confidence levels. This is in agreement with the results about structural identifiability of mLTK model. In fact, $\lambda$ is a structurally non-identifiable parameter for mLTK and, from the theory about model identifiability, we know that structural non-identifiability implies practical non-identifiability.

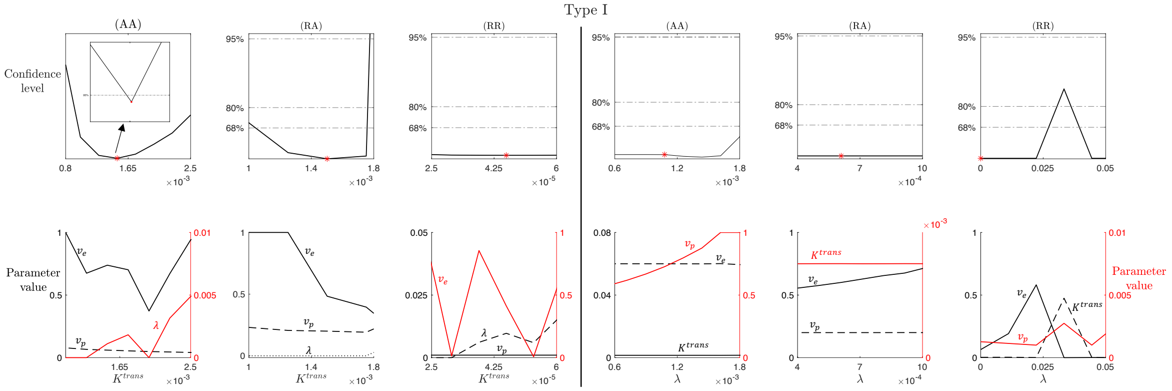


Figure S.3: **$K^{trans}$ and leakage ($\lambda$) practical identifiability for mLTK model and Type I enhancement curve.** Top row: profile likelihood and confidence levels at 68%, 80%, and 95% for the parameter $K^{trans}$ (columns one to three) and $\lambda$ (columns four to six) in the (AA), (RA), and (RR) case for the Type I enhancement curve. Insets in the first and fourth subplot show a zoom of the region around the best fitted value $\hat{K}^{trans}$ and $\hat{\lambda}$ (red markers), respectively. Bottom row: compensating profiles of the parameters $v_e$, $v_p$, and $\lambda$ with respect to variation of $K^{trans}$ around its best fitted value (columns one to three) and of the parameters $K^{trans}$, $v_e$, and $v_p$ with respect to variation of $\lambda$ around its best fitted value (columns four to six). Variation of $\pm 50\%$ around the optimal values of $K^{trans}$ and $\lambda$ are considered.

#### S.3 Supplementary data for the LTK Model

This Section collects additional data and results concerning the analysis of the LTK model proposed in these notes. Table S.1 summarizes the best values for LTK parameter obtained from the data fitting based on a PS algorithm for the three types of CA time-enhancement curves in the three cases of study (AA), (RA), and (RR).

| Case | Type | $K^{trans}$ ($s^{-1}$) | $v_e$ | $v_p$ | $\lambda$ ($s^{-1}$) |
|---|---|---|---|---|---|
| (AA) | I | 0.0022 | 0.6193 | 0.0499 | 0.0104 |
| (AA) | II | 0.009 | 0.5001 | 0.1001 | 0.001 |
| (AA) | III | 0.03 | 0.05 | 0.1 | 0.001 |
| (RA) | I | 0.0025 | 0.4973 | 0.1 | 0.001 |
| (RA) | II | 0.009 | 0.5 | 0.1 | 0.001 |
| (RA) | III | 0.8018 | 0.0326 | 0.8684 | 0 |
| (RR) | I | 0.000449 | 0.0106 | 0.001 | $10^{-6}$ |
| (RR) | II | 0.1396 | 0.2315 | 0.0033 | 0.000482 |
| (RR) | III | 0.1793 | 0.001 | 0.1632 | 0 |

Table S.1: Best parameter values obtained with PS algorithm for the three types of CA time-enhancement curves and in the three cases of study.

Figures S.4 and S.5 collect the results of the practical identifiability analysis for the parameters $K^{trans}$ and $\lambda$ for Type II and III CA time-enhancement curves. In both Figures, Type II and Type III time-enhancement profiles are in line with the ones obtained for Type I in Figures 3 and 4. For the artificial dataset and population-based estimation of the VIF ((AA) case), $K^{trans}$ and $\lambda$ practical identifiability is confirmed with finite confidence region for the 95% confidence level, while it is not possible to define lower/upper (or both) bound of the confidence region for the given confidence levels when an individual-based estimation of the VIF in both (RA) and (RR) cases is considered.

##### S.3.1 RR - Breast cancer tissue

In this Section, we analyze the practical identifiability of the LTK model in breast cancer tissue under the (RR) case. Figure S.6 shows the LTK model fit to the data using Type I, II, and III time-enhancement curves. For breast cancer tissue, the leakage compartment introduced with the LTK and mLTK models calibrates to zero, and is thus not necessary. This indicates that leakage in breast cancer is less important than in GBM. Application to GBM, where large regions of necrosis with slow contrast uptake were the original motivation for

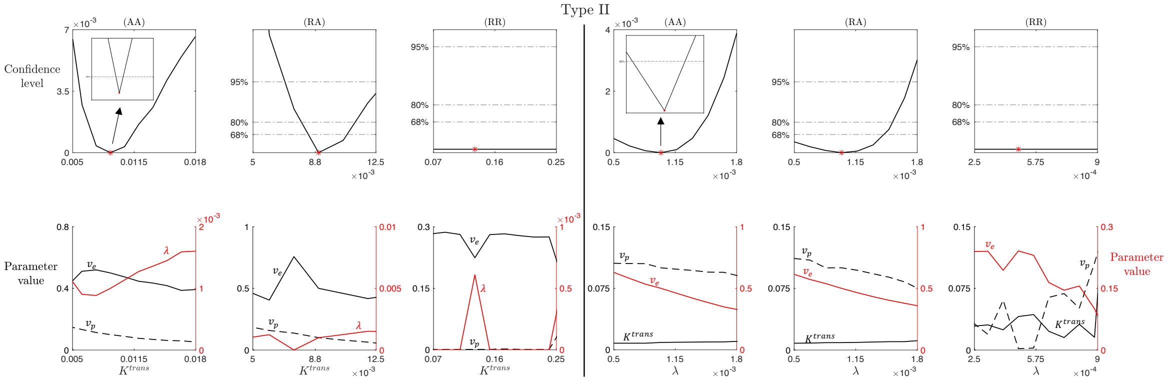


Figure S.4: $K^{trans}$ **and leakage ($\lambda$) practical identifiability for LTK model and Type II enhancement curve.** Top row: profile likelihood and confidence levels at 68%, 80%, and 95% for the parameter $K^{trans}$ (columns one to three) and $\lambda$ (columns four to six) in the (AA), (RA), and (RR) case for the Type II enhancement curve. Insets in the first and fourth subplot show a zoom of the region around the best fitted value $\hat{K}^{trans}$ and $\hat{\lambda}$ (red markers), respectively. Bottom row: compensating profiles of the parameters $v_e$, $v_p$, and $\lambda$ with respect to variation of $K^{trans}$ around its best fitted value (columns one to three) and of the parameters $K^{trans}$, $v_e$, and $v_p$ with respect to variation of $\lambda$ around its best fitted value (columns four to six). Variation of $\pm 50\%$ around the optimal value of $K^{trans}$ and $\lambda$ are considered.

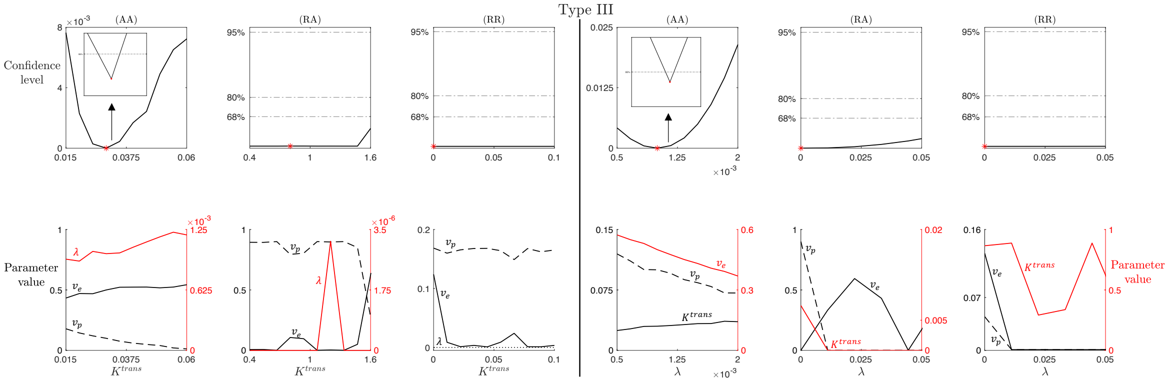


Figure S.5: $K^{trans}$ **and leakage ($\lambda$) practical identifiability for LTK model and Type III enhancement curve.** Top row: profile likelihood and confidence levels at 68%, 80%, and 95% for the parameter $K^{trans}$ (columns one to three) and $\lambda$ (columns four to six) in the (AA), (RA), and (RR) case for the Type III enhancement curve. Insets in the first and fourth subplot show a zoom of the region around the best fitted value $\hat{K}^{trans}$ and $\hat{\lambda}$ (red markers), respectively. Bottom row: compensating profiles of the parameters $v_e$, $v_p$, and $\lambda$ with respect to variation of $K^{trans}$ around its best fitted value (columns one to three) and of the parameter $K^{trans}$, $v_e$, and $v_p$ with respect to variation of $\lambda$ around its best fitted value (columns four to six). Variation of $\pm 50\%$ around the optimal value of $K^{trans}$ and $\lambda$ are considered.

including a leakage compartment [72, 27]. As breast tumors are often highly perfused [73, 74, 75], it follows that leakage often calibrates to 0. Despite this, we analyzed practical identifiability of $K^{trans}$ and $\lambda$, varying the latter in the neighborhood of 0 to observe how the profile likelihood and compensating profiles evolve.

Figure S.7 shows the results about practical identifiability for $K^{trans}$ and $\lambda$ for Type I time-enhancement curve in the (RR) case when breast cancer tissue is considered.
As shown in the last columns of Figures 3 and 4, both $K^{trans}$ and $\lambda$ result practically non-identifiable parameters, independently from the characteristics of the considered tissue. It is not possible, in fact, to define lower and upper bounds for the confidence regions for any of the considered confidence levels.

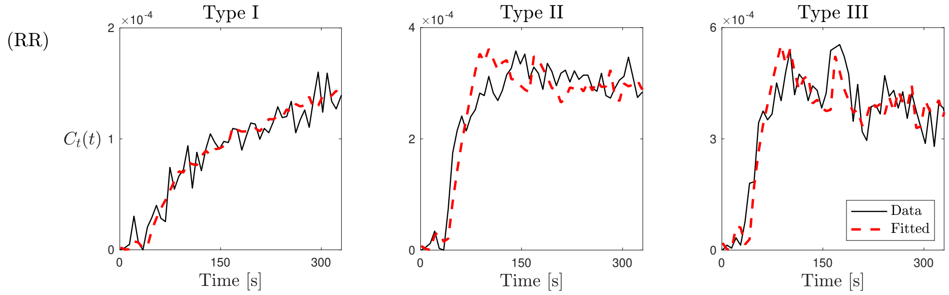


Figure S.6: **Best fitting of the CA evolution with the LTK model** (9)**.** The three types of CA time-enhancement curves (columns) are shown or the (RR) case of the breast cancer tissue.

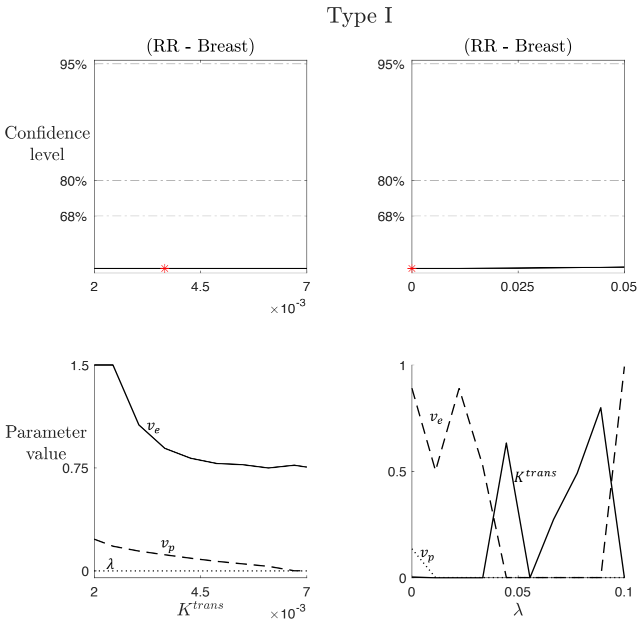


Figure S.7: $K^{trans}$ **and leakage (**$\lambda$**) practical identifiability for LTK model and Type I enhancement curve for breast cancer tissue.** Top row: profile likelihood and confidence levels at 68%, 80%, and 95% for the parameter $K^{trans}$ (first column) and $\lambda$ (second column) in the (RR) case for breast cancer tissue and for the Type I time-enhancement curve. Bottom row: compensating profiles of the parameters $v_e$, $v_p$, and $\lambda$ with respect to variation of $K^{trans}$ around its best fitted value (first column) and of the parameters $K^{trans}$, $v_e$, and $v_p$ with respect to variation of $\lambda$ around its best fitted value (second column). Red markers indicate the best fitted values $\hat{K}^{trans}$ and $\hat{\lambda}$. Variation of $\pm 50\%$ around the optimal value of $K^{trans}$ and $\lambda$ are considered.
